# The interdomain helix between the kinase and RNase domains of IRE1α transmits the conformational change that underlies ER stress-induced activation

**DOI:** 10.1101/2020.01.14.902395

**Authors:** Daniela Ricci, Stephen Tutton, Ilaria Marrocco, Mingjie Ying, Daniel Blumenthal, Daniela Eletto, Jade Vargas, Sarah Boyle, Hossein Fazelinia, James C. Paton, Adrienne W. Paton, Chih-Hang Anthony Tang, Chih-Chi Andrew Hu, Tali Gidalevitz, Yair Argon

**Affiliations:** Division of Cell Pathology, the Children’s Hospital of Philadelphia and the University of Pennsylvania, Civic Center Blvd, Philadelphia, PA 19104; Department of Biology, Drexel University, Philadelphia, PA 19104; Research Centre for Infectious Diseases, Department of Molecular and Biomedical Science, University of Adelaide, Adelaide, SA, 5005 Australia; Wistar Institute, Philadelphia, PA 19104

**Author notes:** University of Salerno, Italy (Daniela Eletto). Department of Biological Regulation, Weizmann Institute of Science, Rehovot, Israel (Ilaria Marrocco).

**Keywords:** Kinase RNase interdomain helix, RNase activity, IRE1 oligomerization, differential autophosphorylation, conformational change

## Abstract

The unfolded protein response (UPR) plays an evolutionarily conserved role in homeostasis, and its dysregulation often leads to human disease, including diabetes and cancer. IRE1α is a major transducer that conveys endoplasmic reticulum (ER) stress to biochemical signals, yet major gaps persist in our understanding of how the detection of stress is converted to one of several molecular outcomes. It is known that upon sensing unfolded proteins via its ER luminal domain, IRE1α dimerizes and oligomerizes (often visualized as clustering), and then trans-autophosphorylates. The IRE1α kinase activity is required for activation of its RNase effector domain and for clustering of IRE1α. It is not yet clear if IRE1α clustering is a platform for the RNase activity, or if the two represent distinct biological functions. Here, we uncover a previously unrecognized role for helix αK between IRE1α kinase and RNase domains in conveying critical conformational changes. Using mutants within this inter-domain helix, we show for the first time that: 1) distinct substitutions (specifically, of Leu827) selectively affect oligomerization, RNase activity, and, unexpectedly, the kinase activity of IRE1α; 2) RNase activation can be uncoupled from IRE1α oligomerization, and phosphorylation of S729 marks the former but not the latter; 3) The nature of residue 827 determines the conformation that the IRE1α protein adopts, leading to different patterns of biochemical activities. In summary, this work reveals a previously unappreciated role for the inter-domain helix as a pivotal conduit for attaining the stress-responsive conformation of IRE1α.

## Introduction

The endoplasmic reticulum (ER) is highly specialized for the folding and quality control of secreted, plasma membrane and organelle proteins. Several conditions, both pathological and physiological, can lead to ER stress, which is characterized by accumulation of misfolded proteins in the ER. This leads to a cellular response called unfolded protein response (UPR), a signal transduction pathway that initiates a transcriptional program to increase the folding capacity of the cell in an attempt to alleviate the stress. If the stress is not resolved, the cell will go through an apoptotic program [1]. ER stress can be chemically induced (e.g. by inhibiting the ER calcium pump with thapsigargin, by inhibiting glycosylation with tunicamycin, or by blocking disulfide bond formation with dithiothreitol, DTT), or can be induced by different physiological processes. In particular, professional secretory cells such as pancreatic β cells and plasma cells have to produce high levels of proteins and experience ER stress [2-6].

It is essential for the cell to sense ER stress promptly in order to initiate the proper coping response. In mammalian cells, sensing ER stress is performed by three different ER transmembrane proteins: Inositol-requiring enzyme 1 α (IRE1α), protein kinase RNA (PKR)-like ER kinase (PERK) and activating transcription factor-6 (ATF6). The IRE1α arm is the most conserved of the three sensors and has been extensively studied in the past. Two mechanisms have been proposed for its activation by ER stress: direct binding to misfolded proteins [7] or the dissociation of the ER luminal chaperone, the binding protein (BiP/GRP78), which keeps IRE1α inactive [8, 9] and sequesters it in this form [10]. IRE1α then dimerizes through its luminal domain [11] and undergoes auto-transphosphorylation via its cytoplasmic kinase domain [12]. This in turn, triggers a conformational change that activates the RNase domains of IRE1α [13]. The enzymatically active IRE1α either cleaves XBP1 RNA at two sites, to create the reading frame for the active XBP1 transcription factor, or cleaves a select group of transcripts once each, initiating degradation of these RNAs (also known as RIDD activity) [14].

This sequence of events that activates IRE1α was defined largely by mutational analysis that showed that mutations in the luminal domain, in the cytosolic kinase domain or in the cytosolic RNase domain can render IRE1α inactive and abrogate the unfolded protein response. How the three domains of the molecule form a modular activation relay is not yet understood. Understanding this mechanism is important because IRE1α can be activated not only by luminal ER stress but also by membrane perturbations that do not require the luminal domain [15], as well as by a bypass mode in response to binding of flavonoids to the cytosolic dimer interface of the RNase domains [16].

It had long been thought that the critical event during mammalian IRE1α activation is the phosphoryl transfer reaction [17]. Counter-intuitively however, some kinase-inactive IRE1α mutants can support RNase activity if provided with appropriate nucleotide mimetics that can bind in the kinase pocket, but are not hydrolyzed [18-20]. This indicates that the key event is not the phosphoryl transfer reaction *per se*, but the conformational change that it triggers. The nature of this conformation change, however, is currently not defined. Yeast IRE1α had been crystalized in two states: an RNase-inactive conformation with the kinase sites of the two monomers oriented ‘face-to-face’, and an RNase ‘active’ back-to-back conformation [17, 21, 22]. If mammalian IRE1α adopts similar orientations, the transition between them could be the conformational change relevant to the ER stress activation. Therefore, it is important to define the intramolecular changes within IRE1α that occur after the kinase activation and initiate RNase activity.

Towards this end, we characterized mutations in helix αK, connecting the kinase and RNase domains, which unexpectedly affect the enzymatic activities of IRE1α dramatically. We show here that some substitutions in helix αK, specifically on residue L827, render IRE1α enzymatically inactive, even though the mutant protein still dimerizes and inhibits a WT version of IRE1α. Other substitutions inhibit the clustering of IRE1α but do not preclude the RNase activities. Importantly, we show that mutations in helix αK have long-range effects on the conformation of IRE1α when it is activated by ER stress, including differential phosphorylation that drives clustering of IRE1α independently of RNase activity [23].

## Materials and Methods

### Cell culture and reagents

HAP1 is a near-haploid human cell line (HZGHC000742c006) derived from the male chronic myelogenous leukemia (CML) cell line KBM-7 (Horizon Discovery, Cambridge, UK). A derivative of HAP1, termed HAP1KO, was engineered by CRISPR/Cas9 editing, to contain a 8-bp deletion in a coding exon of the endogenous IRE1α gene (ERN1), abolishing expression of IRE1α. HAP1 and HAP1KO cells were maintained in IMDM medium (Invitrogen) supplemented with 5% FBS (Atlanta Biologicals), 1% penicillin/streptomycin (Gibco) and 1% sodium pyruvate (Corning, NY). Kms11 is a human myeloma cell line [24]that was grown in RPMI 1640 (Mediatech) medium, supplemented with 10% FBS, 1% penicillin/streptomycin and 1% sodium pyruvate (Sigma, St. Louis).

HAP1, HAP1KO and Kms11 cells were transduced with a Tet-On lentivirus (G418 resistant) and with pTIGHT-IRE1-GFP-HA (IRE1GFP) lentivirus (puromycin resistant) and selected for antibiotic resistance (G418 - 1 mg/ml, puromycin - 2 μg/ml). IRE1GFP contained the human IRE1α sequence and a super-folded GFP grafted in-frame between codons 494 and 495, plus an HA peptide at the C-terminus of IRE1α.

HAP1KO stably expressing the lentivirus pTIGHT-IRE1-GFP-HA (IRE1GFP) (G418 and puromycin resistant) were re-cloned (∼80% transgene expression), and are designated IRE1GFP WT or WT cells. HAP1KO stably expressing IRE1α point mutants were re-cloned similarly. Where indicated, IRE1GFP WT and mutants with only GFP or HA tag were also used. IRE1GFP expression was induced with 1μg/ml doxycycline (dox) (Biochemika) for 16-24 hrs.

Tunicamycin and 4μ8c were purchased from Calbiochem, thapsigargin was from MP Biomedicals and Luteolin was from Sigma. GFP-Trap® A beads were obtained from Chromotek and trypsin from Promega.

### Mutagenesis

The pTIGHT-IRE1-GFP-HA WT plasmid was used as a template for site-directed mutagenesis according to [25]. Pfu Ultra II Fusion HS polymerase was purchased from Agilent. All mutations were validated by Sanger sequencing. Primers used:

L827P: TCAGCGAAGCACGTGGCCAAACACCCGTTCTTC;

P827L: GAAGCACGTGCTCAAACACCCGTTCTT;

L827F:TCAGCGAAGCACGTGTTCAAACACCCGTTCTTC;

L827Q:TCAGCGAAGCACGTGCAGAAACACCCGTTCTTCTG;

L827R:TCAGCGAAGCACGTGCGCAAACACCCGTTCTTC;

D123P: CTCTACATGGGTAAAAAGCAGCCCATCT.

### In silico analysis of protein structure

Rosetta (release 3.11) was used to predict the changes in protein stability due to the point mutations. The input was the crystal structure of apo human IRE1α (PDB: 5HGI). It was pre-minimized using the “minimize_with_cst” application in Rosetta. We followed a ΔΔG _monomer application described by Kellogg et al. [26]for estimating stability changes in monomeric proteins in response to point mutations. In brief, this application uses the input structure of the wild type protein to generate a structural model of the point-mutant. The ΔΔG is given by the difference in Rosetta Energy Unit (REU) between the wild-type structure and the point mutant structure. For more precise calculation, 50 models for each wild-type and mutant structures were generated, and the most accurate ΔΔG was calculated as the difference between the mean of the top three scoring wild type structures and the top three scoring point-mutant structures. The “show_bumps” plugin in PyMol was used to depict the potential steric hinderance and van der Waals clashes in the wild-type and modeled structures.

### RNA extraction, PCR and qPCR

Total RNA was isolated with the Trizol reagent (Life Technologies), following manufacturer’s instructions. 200ng RNA were retrotranscribed to cDNA by priming with oligo(dT)12-18 and Superscript II retrotranscriptase (Invitrogen). Primers to detect human unspliced/spliced XBP1: fwd: AAACAGAGTAGCAGCTCAGACTGC; rev: TCCTTCTGGGTAGACCTCTGGGAG. Quantitative PCR was performed using SYBR green reagent (Applied Biosystems) and the reaction run on Applied Biosystems StepOne Plus machine. Data was analyzed using delta-delta-Ct method. qPCR primers: Rpl19: fwd: AAAACAAGCGGATTCTCATGGA; rev: TGCGTGCTTCCTTGGTCTTAG; Blos1: fwd: CAAGGAGCTGCAGGAGAAGA; rev: GCCTGGTTGAAGTTCTCCAC; CHOP: fwd: GGAGCTGGAAGCCTGGTATG; rev: AAGCAGGGTCAAGAGTGGTG.

### Immunoprecipitation

Cells were lysed in lysis buffer (50mM Tris-HCl pH 8, 150mM NaCl, 5mM KCl, 5mM MgCl2, 1% NP-40, 20mM iodoacetamide, 1X protease inhibitors (Roche)). 5% of the volume of the lysate was saved as “input” in sample buffer and the rest were diluted in TNNB buffer (50mM Tris pH 7.5, 250 mM NaCl, 0.5% NP-40, 0.1% BSA, 0.02% NaN3). GFP-Trap®_A beads were added and incubated for 1 hr at 4° C. After washing, beads were resuspended in sample buffer, boiled for 5 minutes and the proteins were analyzed by SDS-PAGE and western blot.

### Western blots

Cells were lysed in lysis buffer (50mM Tris-HCl pH 8, 150mM NaCl, 5mM KCl, 5mM MgCl_2_, 1% NP-40, 20mM iodoacetamide, 1X protease inhibitors (Roche)). Protein content was determined by BCA assay (Pierce) and proteins were separated by SDS-PAGE and transferred onto nitrocellulose membrane (Bio-Rad). Membranes were blocked, probed with primary and secondary antibodies and scanned on an Odyssey Infrared imager (Li-Cor).

Primary antibodies used: anti-IRE1α: (Cell Signaling Technology); anti-HA: (Covance); anti-14.3.3 (Santa Cruz); anti-phospho-IRE1α Ser724 (Novus Biologicals); anti-phospho-IRE1α Ser729 (Dr. Hu, Wistar Institute, Philadelphia). IRDye-conjugated secondary antibodies were from Li-Cor.

### Limited proteolysis

Cells were lysed in lysis buffer (50mM Tris-HCl pH 8, 150mM NaCl, 5mM KCl, 5mM MgCl_2_, 1% NP-40, 20mM iodoacetamide), protein content was determined as above, trypsin was added at the indicated final concentration and incubated on ice for 30 minutes. The reaction was stopped by adding sample buffer and boiling the samples for 5 minutes. Western blot was then performed as above.

### Microscopy and image analysis

HAP1 cells were plated on 35mm microscopy grade plastic dishes (Ibidi, Germany). Expression of the exogenous IRE1GFP was induced with dox (1 μg/ml for 16-24 hours). The next day the cells were stained with NucBlue® Live ReadyProbes® Reagent (Thermo Fisher), the medium was replaced with phenol red-free IMDM containing ER stressors and the cells were imaged over 8 hours using a Marianas fluorescence microscope equipped with an OKO Lab CO_2_ enclosure on a Zeiss inverted platform, with a 63X Plan-Apochromat 1.4 NA objective. Images were collected using an ORCA-ER camera (Hamamatsu Photonics) using SlideBook V.6 acquisition software. At each time point, a 4.5-6µm Z-stack of images was collected at intervals of 0.5µm. Exposure times varied between 0.1-0.5 sec, depending on sample intensity, unless otherwise specified. In some experiments, cells were imaged using a 63× Plan Apo 1.4 NA objective on an Axiovert 200M (Zeiss) with a spinning disc confocal system (UltraVIEW ERS6, PerkinElmer). Images were collected using an ORCA Flash 4.0 camera (Hamamatsu Photonics) using Volocity V.6.3.1 acquisition software.

### Colony formation assay

HAP1 or HAP1 L827P IRE1GFP cells were plated in 6 well plates (Corning) at 5,000 cells per well. IRE1GFP expression was induced with dox and medium was changed every two days with fresh dox. Cells were stressed with various concentrations of Tm for the indicated times. The drug was then washed out, the plates were incubated for 6 days at 37°C and were then stained with 0.2% (w/v) crystal violet (Sigma) in 2% ethanol for 1 hour at room temperature. The crystals were then dissolved for in 2% SDS (1 hour) and color was quantified at OD570.

### Statistical analyses

To enumerate cells containing clusters, >200 cells were counted per condition in two or more independent experiments. Images were analyzed and quantified using a home-made cluster analyzer for ImageJ [23].

Statistically significant differences between normally distributed populations were evaluated by student’s t test and by non-parametric tests, when the distributions were non-normal.

## Results

### The inter-domain mutation L827P renders IRE1α inactive

We used a clone of HAP1 cells deficient for IRE1α (HAP1KO) as a host for functional complementation of IRE1α and analysis of structure-function relations [23]. HAP1KO cells only activate the XBP1 transcription factor when transduced with active IRE1α [23], such as the previously described doxycycline (Dox)-inducible GFP- and HA-tagged wild type IRE1α (WT IRE1GFP) (Fig. 1A and [23]).

**Figure 1.**
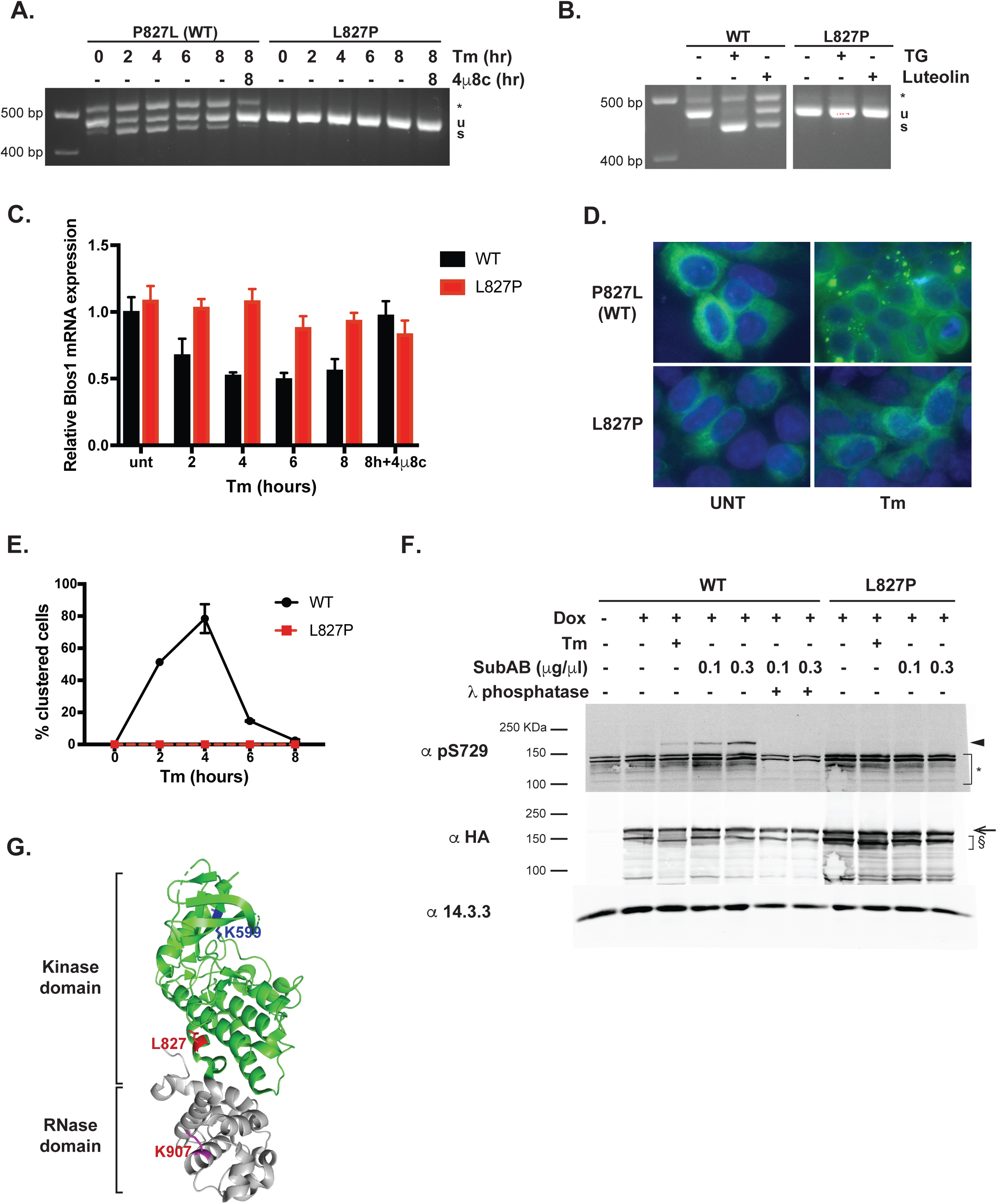
IRE1α L827P is an enzymatically inactive mutant. **A.** Failure of IRE1α L827P to support the XBP1 splicing activity. HAP1KO cells were complemented by either the revertant wild type human IRE1GFP (P827L, see text) or the L827P mutant. Each stable subline was subjected to treatment with 4 μg/ml tunicamycin for the indicated times, after which splicing of XBP1 transcripts was assayed by a RT-PCR gel assay. Specificity of the assay for IRE1α activity was assessed by inclusion of 4μ8c (16 μM) where indicated. u: unspliced XBP1; s: spliced XBP1; *: hybrid band resulting from unspliced and spliced XBP1 (Li et al., 2010). **B.** IRE1GFP L827P cannot be activated by the flavonoid luteolin. HAP1 cells with WT or L827P IRE1α as in A, were treated with 0.2μM thapsigargin, or the IRE1α cytosolic activator luteolin (50 μM), or with the DMSO solvent alone (-) for 2 hr. XBP1 splicing was then assayed as in A. **C.** Failure of IRE1α L827P to support RIDD activity. The same stable sublines were stressed as in A and then the RIDD activity of IRE1α was assayed using qPCR quantitation of the common BLOC1S1 substrate. Data are expressed as the relative abundance of BLOC1S1 under each condition relative to the abundance of the unaffected ribosomal gene Rpl19. Values are means ± SEM of triplicate measurements in two independent experiments. **D.** IRE1GFP L827P does not cluster in response to ER stress. HAP1KO cells re-complemented with WT or L827P IRE1GFP were induced with dox for 16 hr. The following day they were treated with Tm (4 μg/ml) and imaged over 8 hr. Images are representative fields of the 4-hr treatment. **E.** Quantification of clustering. The images from Fig. 1D were quantified and plotted. **F.** L827P IRE1GFP is not phosphorylated on S729 following induction of ER stress. HAP1KO re-complemented with WT or L827P IRE1GFP were stressed with Tm as in D and with SubAB at the indicated concentrations for 2 hr. Cells were lysed and activation of IRE1α was assessed by western blot with an antibody against phosphorylated Ser729 of IRE1α. 14.3.3: housekeeping protein. Arrow indicates full length IRE1GFP; arrowhead: phospho-IRE1GFP S729; §: lower molecular weight bands that appear to be IRE1α specific and size-sensitive to Tm treatment; *: non-specific bands. **G.** Location of L827 in the crystal structure of IRE1α. Residue L827 of human IRE1α is highlighted in red in PDB 5HGI. For orientation, the catalytic residue in the kinase domain, K599, is marked in blue and the catalytic residue in the RNase domain, K907, is marked in purple.

Using this system, the substitution of Leucine 827 with Proline was discovered fortuitously in an IRE1α cDNA that failed to restore XBP1 splicing in response to either Tm (Fig. 1A) or Tg (Fig. 1B). When P827 was mutated back to Leu, the splicing activity was restored in response to Tm-induced ER stress (Fig. 1A, denoted P827L), showing that the inactivation of IRE1α was due to the L827P substitution. The re-cloned revertant protein is referred to hereafter as WT. In addition to its inability to support XBP1 splicing, L827P also fails to perform the RIDD activity of IRE1α, as measured with the BLOC1S1 transcript (Fig. 1C). Thus, the mutant protein is unable to perform either of the RNase activities.

L827P IRE1α is not only refractive in response to ER stress conditions, but also to treatment with the flavonoid luteolin (Fig. 1B), which binds to the RNase domain of IRE1α, enhances dimerization and activates WT IRE1α even without ER stress [16].

A fourth phenotype of L827P IRE1GFP is that it fails to cluster under ER stress (Fig. 1D-E). ER-stress induced clustering is a behavior displayed by both the enzymatically active IRE1α and by IRE1α that is RNase-deficient or inhibited. We define clustering as the switch from reticular dispersion to foci localization of IRE1α molecules in response to ER stress [23]. It has been shown before that WT IRE1GFP clusters in response to Tm in transient fashion over an 8 hour time frame [23, 27-29], but the L827P IRE1GFP molecule did not show any clustering (Fig. 1D-E).

All the activities that we found to be defective in the L827P mutant should depend on the auto-phosphorylation activity of IRE1α [23, 30, 31]. Therefore, we tested the phosphorylation status of L827P IRE1α using an antibody that detects phospho-Ser729 [31]. Phosphorylation of wild type IRE1α was detected after treatment with Tm, and more robust phosphorylation was evident upon ablation of BiP with the toxin SubAB, a treatment known to provide higher level of ER stress (Fig. 1F; [32]). In contrast, no significant phospho-Ser729 was detected even at the high concentration of SubAB treatment for L827P IRE1α (Fig. 1F). Because of the specificity of the antibody, it is possible that L827P is phosphorylated elsewhere, but not on Ser729. Therefore we subjected ER stressed cell lysates to lambda phosphatase and resolved them by electrophoresis to visualize the migration shifts. As shown in Suppl. Fig. 1, Tm -treated or SubAB-treated IRE1α L827P exhibited no significant mobility shifts compared to unstressed IRE1α, indicating that the L827P protein is indeed not phosphorylated upon ER stress. We conclude that L827P specifies a form of IRE1α that lacks most of the activities of the protein even though the mutation is spatially distant from IRE1α catalytically active residues (Fig. 1G).

### L827P IRE1α inhibits wild type IRE1α in leukemic HAP1 and in multiple myeloma Kms11 cells

We next asked if L827P could influence the activity of wild-type IRE1α. We expressed doxycycline-inducible L827P IRE1α in the parental HAP1 cell line that had intact endogenous IRE1α. Prior to induction of the mutant protein these cells exhibited normal XBP1 splicing when treated with either Tg or Tm (Fig. 2A-B, respectively). However, when L827P IRE1GFP expression was induced, the XBP1 splicing activity of the WT allele was progressively inhibited in proportion to the level of induction of L827P IRE1GFP (Fig. 2A-B, E).

**Figure 2.**
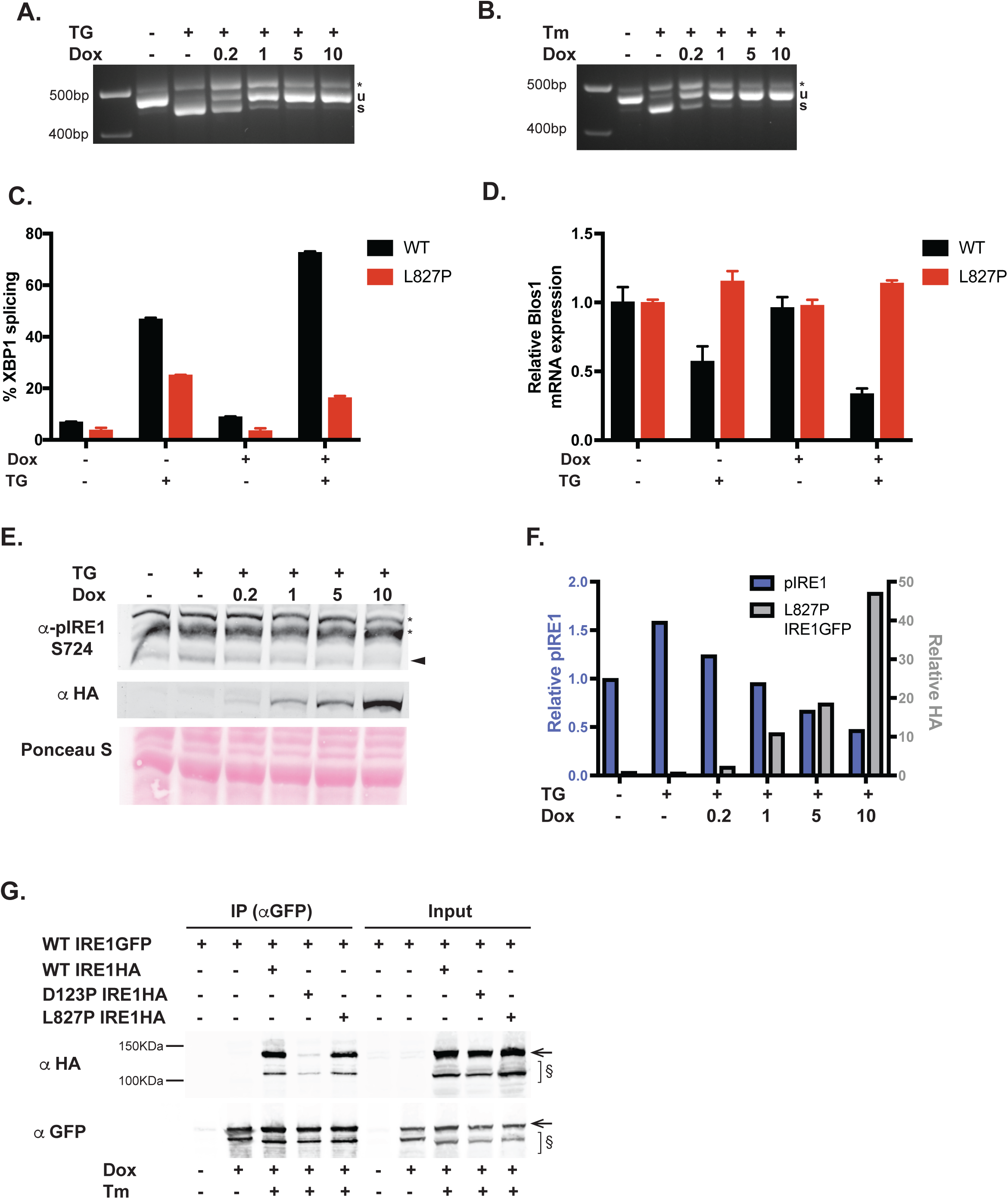
The L827P mutant inhibits WT IRE1α in HAP1 and Kms11 cells. **A.** IRE1GFP L827P inhibits XBP1 splicing in response to thapsigargin in leukemic HAP1 cells. Parental HAP1 cells expressing IRE1GFP L827P in addition to the endogenous WT IRE1α were induced with the indicated concentrations of dox (μg/ml) for 16 hr. The cells were then treated with 0.2 μM Tg for 4 hr and XBP1 splicing was assessed by RT-PCR. **B.** L827P IRE1GFP inhibits XBP1 splicing in response to tunicamycin in leukemic HAP1 cells. The same experiment as in A was performed except the cells were treated with 4 μg/ml Tm for 4 hr. **C.** L827P IRE1GFP inhibits ER stress-induced XBP1 splicing in multiple myeloma Kms11 cells. Kms11 cells expressing WT or L827P IRE1GFP were induced with dox for 16 hr and treated with 0.5 μM Tg for 4 hr where indicated. RNA was extracted and XBP1 splicing was assessed by RT-PCR and quantified. **D.** L827P inhibits RIDD activity in response to ER stress in Kms11 cells. The same samples as in C were used to perform a qPCR to detect BLOC1S1 expression levels, normalized over the unaffected ribosomal gene Rpl19. **E.** Expression of L827P decreases endogenous WT IRE1α phosphorylation in response to ER stress. Parental HAP1 cells, with constitutive expression of WT IRE1α and inducible expression of IRE1GFP L827P were induced with dox as above and then treated with Tg (0.2 μM for 4 hr). Cells were lysed and proteins were analyzed by western blot. Arrowhead: endogenous phospho-S724 IRE1α; *: non-specific bands. **F.** Quantification of phospho-IRE1α S724 and L827 mutant. The intensities of the western blot bands from the experiment described in E were normalized to total protein contents of the samples, measured by Ponceau S staining, quantified and plotted. **G.** L827P binds full-length WT IRE1α. HAP1KO cells were re-complemented with WT IRE1GFP, WT IRE1HA, D123P IRE1HA, L827P IRE1HA or combinations of constructs. The cells were induced with dox for 16 hr and treated with 4 μg/ml Tm for 4 hr where indicated. Cells were collected, lysed and subjected to immunoprecipitation with GFP-Trap beads. Beads-bound proteins were analyzed by western blot. Input: 5% of the lysates. Arrow: full length IRE1GFP or IRE1HA; §: lower molecular weight bands that appear to be IRE1α specific and size-sensitive to Tm treatment.

To test if this inhibitory effect was cell-type specific, we expressed the same doxycycline-inducible WT or L827P IRE1GFP in multiple myeloma Kms11 cells, that express intact endogenous IRE1α. As shown in Suppl. Fig. 2, the exogenous IRE1GFP was expressed at a basal level even in absence of dox, and its level then increased upon addition of dox. Basal level of XBP1 splicing was already lower at steady state in L827P IRE1α cells, in comparison to the WT IRE1α-expressing cells. After ER stress was induced with Tg for 4 hours, L827P IRE1α inhibited endogenous IRE1α activity even further, showing only ∼16% XBP1 splicing in respect to the WT IRE1GFP-transduced cells (Fig. 2C).

A similar inhibition was observed for the RIDD activity. In the presence of L827P, endogenous IRE1α was unable to cleave Blos1 in response to Tg treatment, whereas WT IRE1GFP showed about 60% reduction of Blos1 mRNA (Fig. 2D). Furthermore, co-expression of L827P IRE1GFP, which does not auto-phosphorylate (Fig. 1F, Suppl. Fig. 1), inhibited the phosphorylation of the endogenous WT IRE1α in a dose-dependent manner (Fig. 2E, F). Importantly, dox dose-dependent inhibition of the endogenous IRE1α reflected increasing levels of the mutant protein (Fig. 2E, F). Thus, the mutant IRE1α inactivates the endogenous WT IRE1α, suggesting a dominant-negative interaction. This inhibition is selective for WT IRE1α, as it does not extend to the homologous UPR sensor PERK, as shown by insensitivity of CHOP transcription to the expression of L827P (Suppl. Fig. 3).

To confirm the dominant-negative mechanism, we asked whether L827P mutant protein inhibits the wild type IRE1α by direct interaction. We created two different versions of exogenous IRE1α: tagged with GFP (IRE1GFP) or with HA alone (IRE1HA). We then co-expressed L827P IRE1HA and assessed its association with WT IRE1GFP, via immune-isolation with anti-GFP beads. As shown in Fig. 2G, association of L827P IRE1α with WT IRE1α happens to the same extent as association of two WT monomers. By comparison, the dimerization-impaired D123P IRE1α mutant associated with WT IRE1α much less efficiently. Thus, the L827P mutant protein likely inhibits the wild-type IRE1α by forming mixed dimers.

The self-association of IRE1α monomers is known to be mediated by the luminal domains as well as the trans-membrane segments and the cytoplasmic domains [11, 15]. The inability of the wild-type endogenous protein to phosphorylate the mutant protein suggests that the kinase domains in the mixed dimers are either not within the required interaction distance, or not oriented correctly for trans-phosphorylation of the mutant by wild-type.

### L827P IRE1α inhibits clustering of wild type IRE1α

Since the clustering of IRE1GFP is distinct from the RNase enzymatic activity [23], and since L827P IRE1GFP dimerizes with the WT protein (Fig. 2), we asked whether the two proteins co-cluster when expressed in the same cells, or whether the mutant protein affects the clustering of the WT IRE1α. Towards this end, we replaced the GFP moiety in WT IRE1GFP with mCherry and placed the red fluorescent protein version under a different selection (blasticidin), to allow simultaneous expression of mCherry-tagged WT IRE1α with GFP-tagged L827P IRE1α. We confirmed that IRE1mCherry protein was as responsive to Tm stress as IRE1GFP (Table 1, Fig. 3A and [23]). Cells co-expressing the GFP- and mCherry-tagged WT IRE1α fluorescent proteins had either red, green or yellow reticular pattern, because of the heterogeneous expression levels of each vector (Fig. 3A). Importantly, upon Tm stress, cells expressing both WT proteins formed only yellow clusters, showing that GFP- and mCherry-tagged WT IRE1α proteins interact with each other as efficiently as the singly-tagged proteins (Fig. 3A).

**Figure 3.**
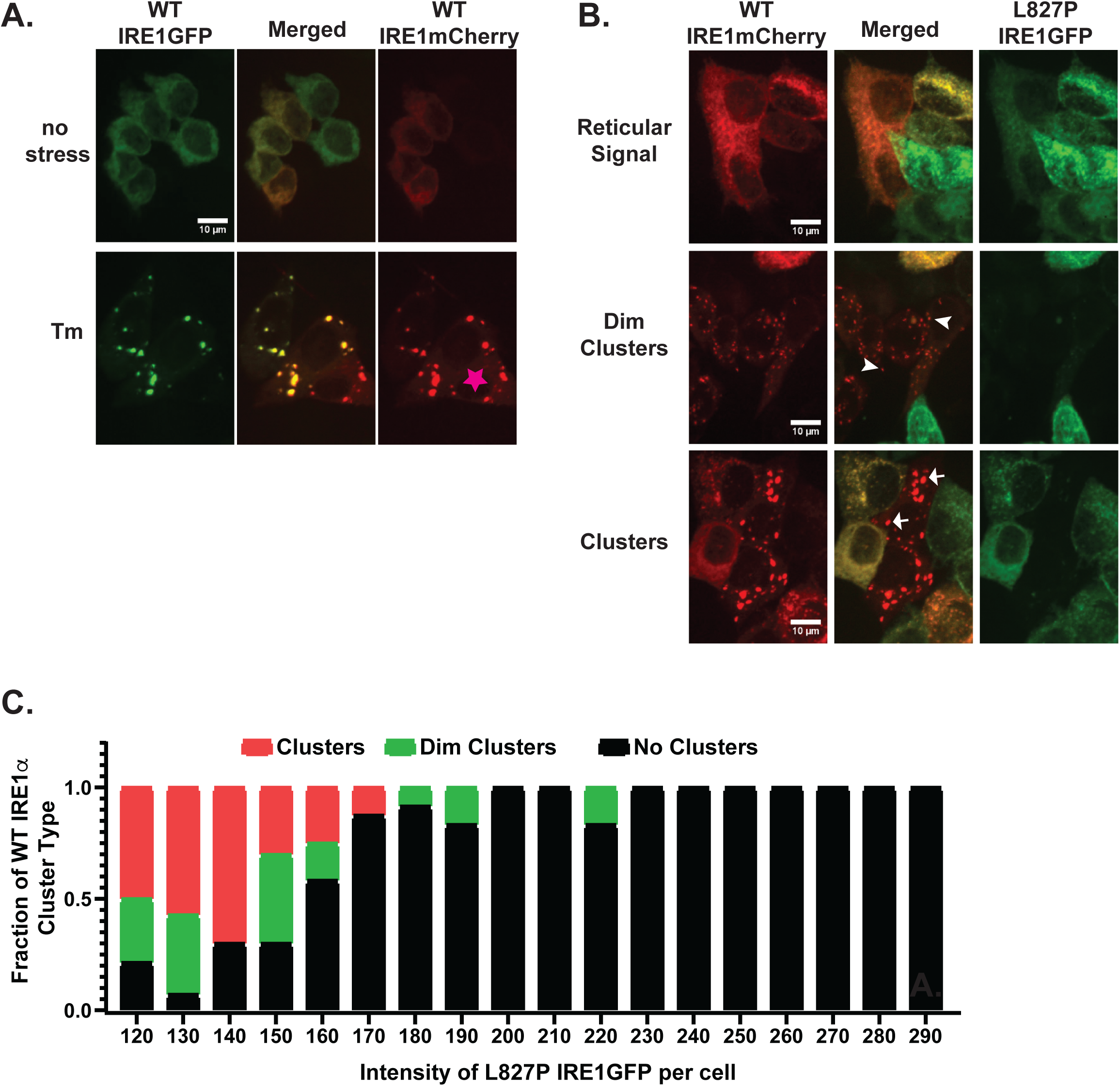
Clustering of WT IRE1α with L827P mutant. **A.** mCherry-tagged IRE1α clusters similarly to IRE1GFP. HAP1KO expressing both WT IRE1GFP and WT IRE1mCherry were stressed with 4 μg/ml Tm for 3hrs and imaged. Representative images are shown. Star, a cell expressing only IRE1mCherry. **B.** Co-expression of mutant L827P IRE1GFP and WT IRE1mCherry. Representative images of WT IRE1mCherry/L827P IRE1GFP-expressing cells treated with 4 μg/ml Tm for 3 hrs. Cells which co-express mCherry and GFP typically exhibit reticular signal with no clusters (reticular signal). mCherry-expressing cells with low GFP expression form faint cluster-like foci termed dim clusters or the bright foci typical of WT IRE1α clusters. Arrowheads: two typical dim clusters; arrows: two typical bright clusters. **C.** L827P inhibits WT IRE1α clustering. WT IRE1mCherry/ L827P IRE1GFP-expressing cells were categorized by cluster type and binned according to the intensity of GFP signal of individual cells.

**Table 1.** Co-clustering of WT and mutant IRE1α. HAP1KO cells expressing a combination of WT or L827P IRE1GFP or IRE1mCherry were treated with 4 μg/ml Tm for 3 hrs, images were taken and cells were scored for different phenotypes. Cells expressing WT IRE1 only, either GFP or mCherry-tagged, clustered in 70-80% of the cases (e.g. Fig. 1E). In presence of the L827P mutation, given the heterogeneity of the protein expression and depending if the levels of the mutant were high and detectable (Red and Green cells column) or not microscopically visible (Red Cells column), WT IRE1α did not cluster or clustering was severely impaired (∼40%), respectively.

|  |  | Red and Green Cells |  |  | Red Cells |  |  |
| --- | --- | --- | --- | --- | --- | --- | --- |
|  | Treatment | # Red cells | # Red Cells with Clusters | % Red Cells with Clusters | # Red cells | # Red Cells with Clusters | % Red Cells with Clusters |
| WT-GFP / WT-mCherry | mock | 57 | 0 | 0.0 | 18 | 0 | 0.0 |
| WT-GFP / WT-mCherry | Tm | 92 | 67 | 72.8 | 33 | 25 | 75.8 |
| L827P-GFP / WT-mCherry | mock | 85 | 0 | 0.0 | 26 | 0 | 0.0 |
| L827P-GFP / WT-mCherry | Tm | 185 | 0 | 0.0 | 83 | 32 | 38.6 |

When WT IRE1mCherry was co-expressed with mutant L827P IRE1GFP, most cells failed to exhibit clusters (Table 1 and Fig. 3B, C). Notably, on the background of bright L827P IRE1GFP, there was complete lack of red WT IRE1mCherry clustering, and only cells with low L827P levels displayed clusters of WT IRE1mCherry, that were usually dim (Fig. 3B, C). This indicates that the presence of 827P IRE1α in the same cell inhibits clustering of WT molecules in proportion to the stoichiometry of the two. Interestingly, a small number of cells contained clusters that were positive for both WT IRE1mCherry and L827P IRE1GFP, but all of these were the atypical dimmer clusters. This suggests that mixed dimers are unable to cluster efficiently, and may indeed interfere with further recruitment of the WT protein. Therefore, L827P IRE1GFP is dominant-negative not only with respect to kinase and RNase activities, but also with respect to stress-induced oligomerization of IRE1α.

### Other L827 substitutions provide a graded loss-of-function phenotype

The L827P mutation is located at the interface between the kinase domain and the RNase domain of IRE1α (Fig. 1G). L827 is within helix αK (S821 to P830 in PDB 4PL3) and is relatively solvent exposed (Fig. 4A). Furthermore, L827 is nestled against Thr674 from helix αE (Fig. 4A) in a relatively hydrophobic surface (residues 665-680 of murine IRE1α (PDB 4PL3) [33]; residues 664-681 of human IRE1α (PDB 5HGI; [34]). The conformation of Leu827 and the interactions between helix αK and its neighboring helices αE and αJ are similar in crystals of the murine or human IRE1α, and are unchanged by binding of nucleotides or inhibitors to the kinase domain or by binding of flavonoids to the RNase domain. Thus, the reasons for the loss-of-function phenotype are not immediately apparent from inspection of the various crystal structures.

**Figure 4.**
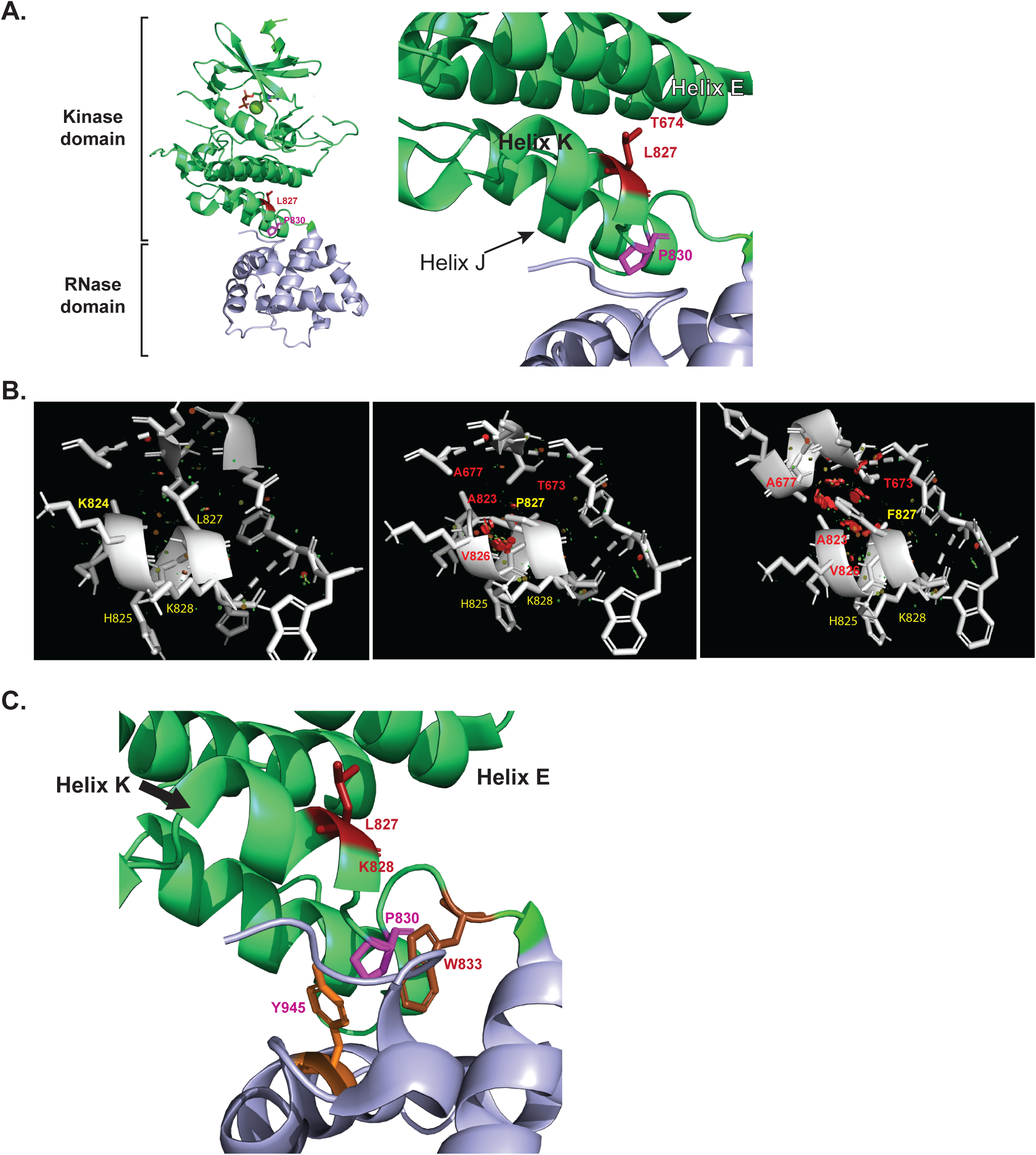
Structural elements of the interdomain helix αK. **A.** Crystal structure of the cytoplasmic portion of human IRE1α (from PDB 5HGI). Kinase domain residues are colored green and RNase domain residues are colored light blue. Left panel, overall structure with residues L827 and P830, as well as the catalytic kinase residue K599 depicted in stick format and highlighted red. Right panel, close-up view of the interdomain helix and L827, which projects from helix αK towards T674 in helix αE. P830 is proximal to the RNase domain. **B.** Predicted intramolecular clashes when Leu827 is mutated to either Pro827 or Phe827. Wild type IRE1α (Leu827, left panel) was substituted *in silico* with Pro827 (center panel) or Phe827 (right panel) and the amino acids within 5Å are depicted. In each case, neighboring residues with predicted molecular clashes (Rosetta) are shown in red, while neighbors without predicted clashes are colored yellow. **C.** Close-up projection of the interdomain helix αK in 5HGI where all the residues whose mutations are described in this work and in [36] are annotated with their side chains in PyMol stick format.

To help explain the profound loss-of-function of L827P we first investigated how substitutions of residue 827 affect the stability of IRE1α: we modeled the cytosolic portion of IRE1α using two algorithms, Rosetta [26] and SDM [35], that predict the changes in protein stability upon substitutions of single amino acids. The Rosetta predicted structural changes, based on the 5HGI crystal structure of human IRE1α, showing that Pro is the most destabilizing substitution at position 827, with an estimated ΔΔG at −2.45 units (Table 2). The substitution of Pro for Leu827 is very likely to cause perturbation in the local conformation of neighboring amino acids, as mapping Pro827 on the structure of human (PDB 5HGI) or murine (PDB 4PL3 or 4PL5) IRE1α suggests that this substitution would clash with Val826, Lys824 and Ala823, the residues that form the turn connecting helices αJ and αK (Fig. 4B). This would weaken the interactions of helices αK-αE and change the geometry of helix αK, at least with respect to Val826, His825 and Lys824. It is quite possible that these main chain alterations could propagate to the RNase active site (Fig. 4A and see more below).

**Table 2.** Predicted stability of IRE1α mutants. Substitutions of Leu827 characterized in this work and the P830 mutations characterized in [36] were modelled in the 5HGI PBD structures of human IRE1α, and the predicted free energy changes calculated according to the Rosetta algorithm [26]. For comparison, many mutations were also modelled in the 4PL5 PDB structure of murine IRE1α and their predicted free energy changes were calculated according to the SDM algorithm [35]. ND, not determined. The destabilization energies of the RNase active site (K907A) and the kinase active site (K599A) also serve as useful comparisons.

| **Mutation** | **Rosetta**<br>**(Energy Units)** | **SDM $\Delta\Delta G$** |
| --- | --- | --- |
| L827P | ND | -2.23 |
| L827P | 234.6 | -2.75 |
| L827I | 3.0 | -0.45 |
| L827V | 4.7 | -2.93 |
| L827F | 109.1 | -1.14 |
| L827Y | 6.9 | -0.93 |
| L827R | 6.3 | -1.35 |
| L827Q | 6.1 | -0.29 |
| L827E | 7.7 | -0.21 |
| L827A | 5.7 | -2.69 |
| P830L | 40.0 | ND |
| P830A | 3.8 | ND |
| K907A | ND | -0.56 |
| K599A | ND | -0.38 |

The Rosetta algorithm also predicts that Phe827, in addition to Pro827, is a destabilizing substitution. All other substitutions are far less destabilizing (Table 2); for example, substituting Leu827 with the aliphatic residues Ile or Val is predicted to have a modest effect, similar to substitutions with the charged residues Arg, Lys, Glu or Asp (Table 2). Modeling Phe827 suggests that 3/4 of the common rotamers would clash with Thr673 in helix αE or with Asp824 and Val826 in helix αK. The Phe rotamer with the fewest clashes is shown in Fig. 4B.

### Arg, Gln and Phe substitutions at position 827 affect IRE1α activities differentially

To test some of these predictions, we generated additional IRE1α mutants with substitutions with Arg, Gln, and Phe at residue 827. When assayed in cells, L827F IRE1GFP is expressed at a similar level to WT IRE1α and as small, variable foci (Fig. 5A, C). Interestingly, this substitution had divergent effect on the different IRE1α activities: L827F foci do not change under Tm stress, resembling the unchanged distribution of the L827P protein (Fig. 5C), even though it splices XBP1 (Figs. 5B, 6A) and performs RIDD (Fig. 6B) as efficiently as the WT protein. Thus, L827F inhibits the clustering phenotype selectively. Most surprisingly, the RNase activity of this mutant is induced without obvious phosphorylation on Ser729 in the kinase activation loop (Fig. 6C). The activation of L827F RNase without phosphorylation on Ser729 is seen in response to either Tm or the even stronger stress of SubAB (Fig. 6C) and therefore, we examined the general phosphorylation status of IRE1α using λ phosphatase-induced electrophoretic shift, as in Suppl. Fig. 1 above. L827F after Tm or SubAB stress resolves into two bands, and at least the larger one is shifted by phosphatase treatment (Suppl. Fig. 1), indicating that it is phosphorylated even though it is not reactive with anti-pSer729. We conclude that L827F is phosphorylated in response to ER stress, but at different positions from wild type IRE1α.

**Figure 5.**
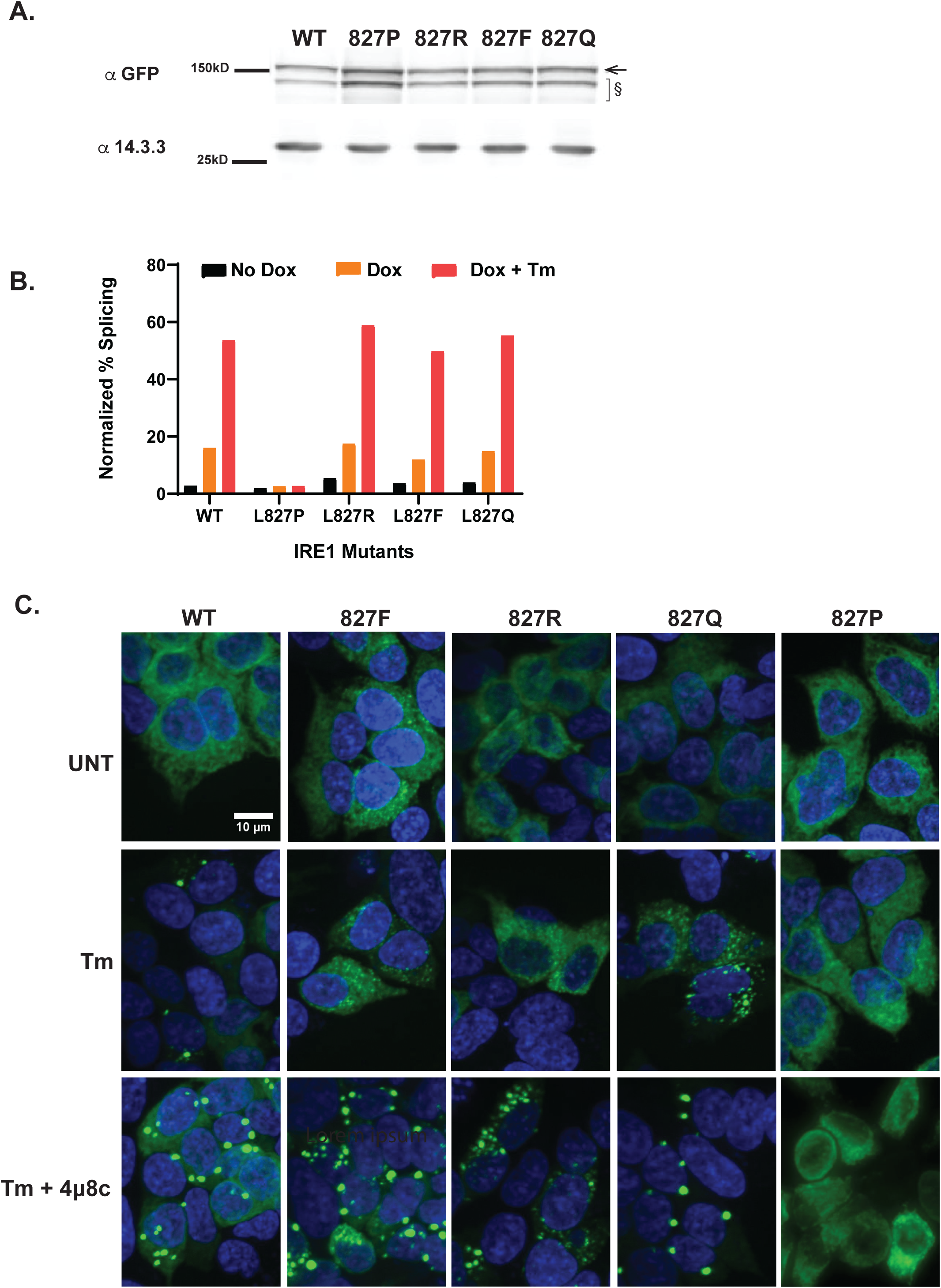
Graded RNase activity and clustering responses of substitutions of L827. **A.** Expression of L827 mutants. HAP1KO cells were infected with lentiviruses expressing the indicated IRE1GFP constructs and selected by puromycin to obtain stable clones. Detergent lysates of each were resolved by SDS-PAGE and probed with anti-GFP. Anti-14.3.3 probing served as a loading control. Arrow: full length IRE1GFP; §: lower molecular weight bands that appear to be IRE1α specific. **B.** RNase activity of L827 substitutions. IRE1α expression was induced in the stable clones with dox and the cells were stressed by treatment with 4 μg/ml Tm for 4 hr to activate XBP1 splicing where indicated. The percentage of splicing was measured from gel analysis and normalized to the protein expression as measured in A. **C.** Clustering behavior of L827 substitutions. Representative fields of stable clones that were subjected to ER stress (Tm, 4 μg/ml) for 4 hrs, or left untreated (UNT) and imaged. A combined treatment of Tm plus 4μ8c (16 μM) was used to generate the mega-clusters, as described by Ricci et al. [23].

**Figure 6.**
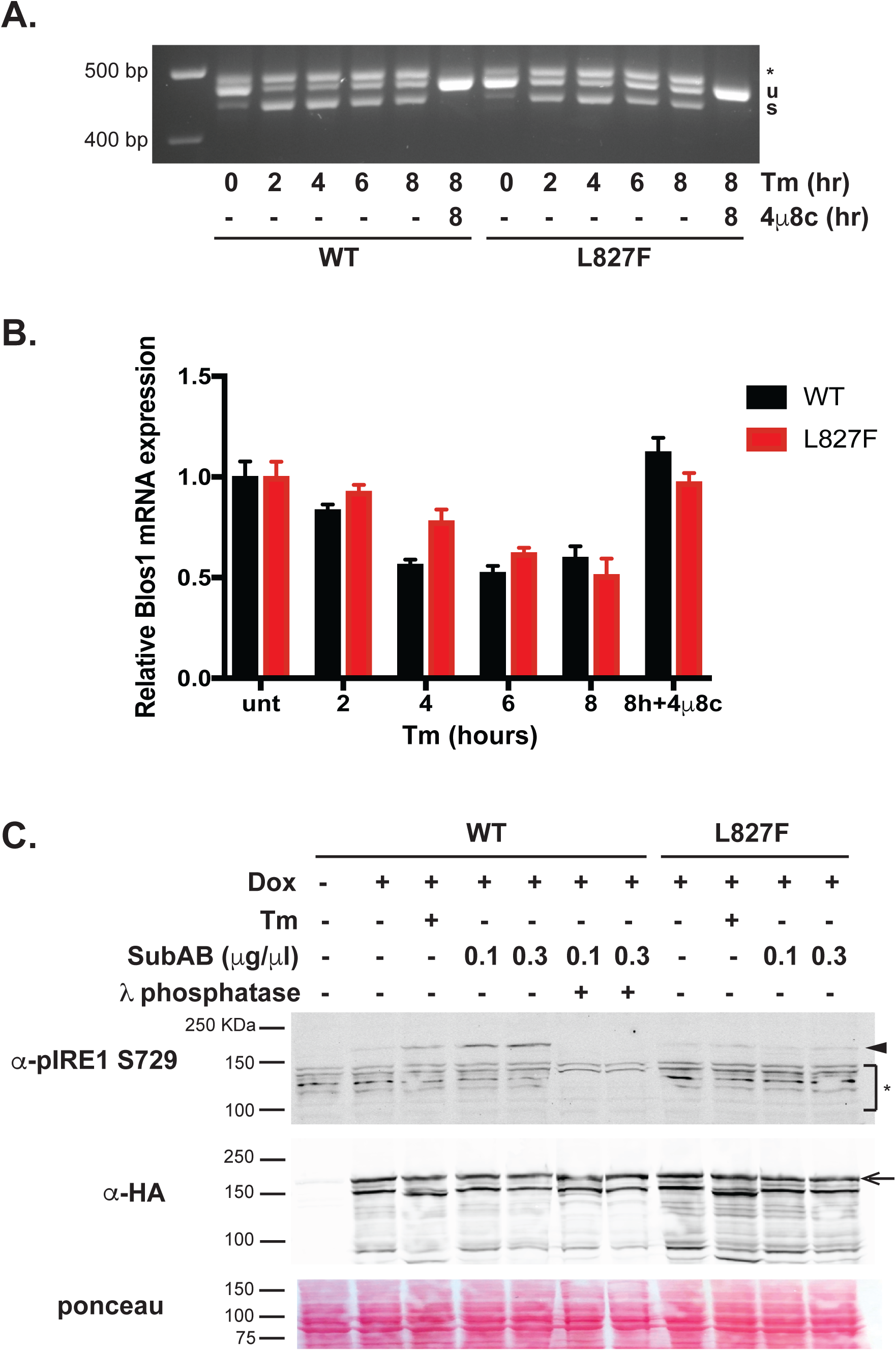
L827F phosphorylation but not RNase activity differs from WT IRE1α. **A.** L827F IRE1α can perform XBP1 splicing activity. HAP1 KO IRE1GFP WT or L827F cells were treated with 4 μg/ml Tm or 16 μM 4μ8c for the indicated times, RNA was extracted, XBP1 splicing assessed by RT-PCR and run on agarose gel. Band identity is as in Fig. 1A. **B.** L827F IRE1α is capable of cleaving RIDD substrates. HAP1 KO IRE1GFP WT or L827F cells were treated as in A, then the RIDD activity of IRE1α was assayed using qPCR quantitation of the common BLOC1S1 substrate. Data are expressed as the relative abundance of BLOC1S1 under each condition relative to the abundance of the unaffected ribosomal gene Rpl19. Values are means ± SEM of triplicate measurements in two independent experiments. **C.** L827F IRE1GFP is not phosphorylated on S729 following induction of ER stress. HAP1KO cells re-complemented with WT or L827F IRE1GFP were stressed with Tm (4 μg/ml) or SubAB at the indicated concentrations, for 2 hr. Cells were lysed and activation of IRE1α was assessed by probing the blots first with an antibody against phosphorylated Ser729 and re-probing with anti-HA tag. Arrow: full length IRE1GFP; arrowhead: phospho-S729 IRE1GFP; *: aspecific bands.

In addition to L827F, substitution of Leu827 with Glutamine also enables XBP1 splicing activity (Fig. 5B), but unlike L827F it partly restores clustering (Fig. 5C). The L827Q clusters were similar in size and number per cell to the WT clusters but the L827Q clusters are duller than WT IRE1α ones and a smaller fraction of the total IRE1α is incorporated into them (more IRE1α remains reticular) (Fig. 5C and Suppl. Fig 4A). These observations are broadly in line with the free energy predictions and suggest that L827Q molecules are active, but do not pack as efficiently as WT into clusters. All these aspects suggest that the observed protein structural perturbations are graded as predicted by the modelling algorithms.

It has been previously shown that a mutation in the kinase-RNase interface, P830L, renders IRE1α more unstable [36]. However, all of the mutants we tested showed a similar half-life to WT IRE1α (Suppl. Fig. 5), suggesting that the observed effects on IRE1α activities are not due to protein stability.

As we showed previously [23], enzymatically inactive IRE1α can still cluster, and in fact forms larger and more persistent clusters. Therefore, we asked whether mutants that are deficient in clustering in response to ER stress can still cluster under the stress when inactivated with the inhibitor 4μ8c. L827Q, L827R and L827F respond to 4μ8c+Tm and form aggregates that are indistinguishable from the large clusters formed by WT IRE1α when inhibited (Fig. 5C). Thus, the substitutions L827F, L827R and L827Q (but not L827P) are inherently competent to cluster, but are defective in their response to Tm stress.

We conclude that RNase activity and oligomerization are distinct and respond differently to alterations in the helix αK element; some substitutions preserve the RNase activity but still impair stress-induced oligomerization of IRE1α.

### L827P and L827F have conformations distinct from wild-type IRE1α

Our prediction anticipated a perturbation in the local conformation in the area around the L827P mutation but the effects of the mutation are observed in IRE1α enzymatic activities carried out in distal residues (kinase and RNase domains). For these reasons, we asked whether other substitutions of helix αK residues lead the IRE1α molecule to assume distinct conformations from the wild type. In order to assess that we performed limited trypsin proteolysis experiments. Lysates of HAP1KO IRE1GFP WT or L827P, either untreated or stressed with 4 μg/ml Tm for 4 hours were subjected to low dose trypsin (0.25-10 μg/ml) digestion. The resolved proteins were analyzed by western blot using two different antibodies: anti-HA that recognizes a tag at the C-terminus of IRE1α and anti-GFP, located in the same molecule between the transmembrane and the kinase domains. The analysis revealed the presence of unique tryptic fragments that characterize the WT and the L827P mutant (Fig. 7A), indicating that the two molecules are differentially susceptible to trypsin treatment. For example, the ∼55kD fragment containing HA which is more prevalent in the L827P mutant (Fig. 7A) indicates a more accessible tryptic site in the N-terminal lobe of the kinase domain. Other informative fragments are exemplified by the doublet at 36kD which is more resistant to trypsin when expressed in the WT lysate of unstressed cells and more sensitive in the L827P lysate (Fig. 7A). The size of these peptides shows that sequence alterations at residue 827 affect the conformation at distant amino acids. The numbers of affected peptides allow the conclusion that distinct sites are more exposed to trypsin in L827P than in WT IRE1α under ER stress. Remarkably, when L827F was subjected to similar partial proteolysis, the proteolysis pattern resembled that of the mutant L827P and not the WT protein, despite the phenotype (Fig. 7B). Thus, the trypsin sensitivity assay reflects the phosphorylation and clustering activities more than the RNase activity.

**Figure 7.**
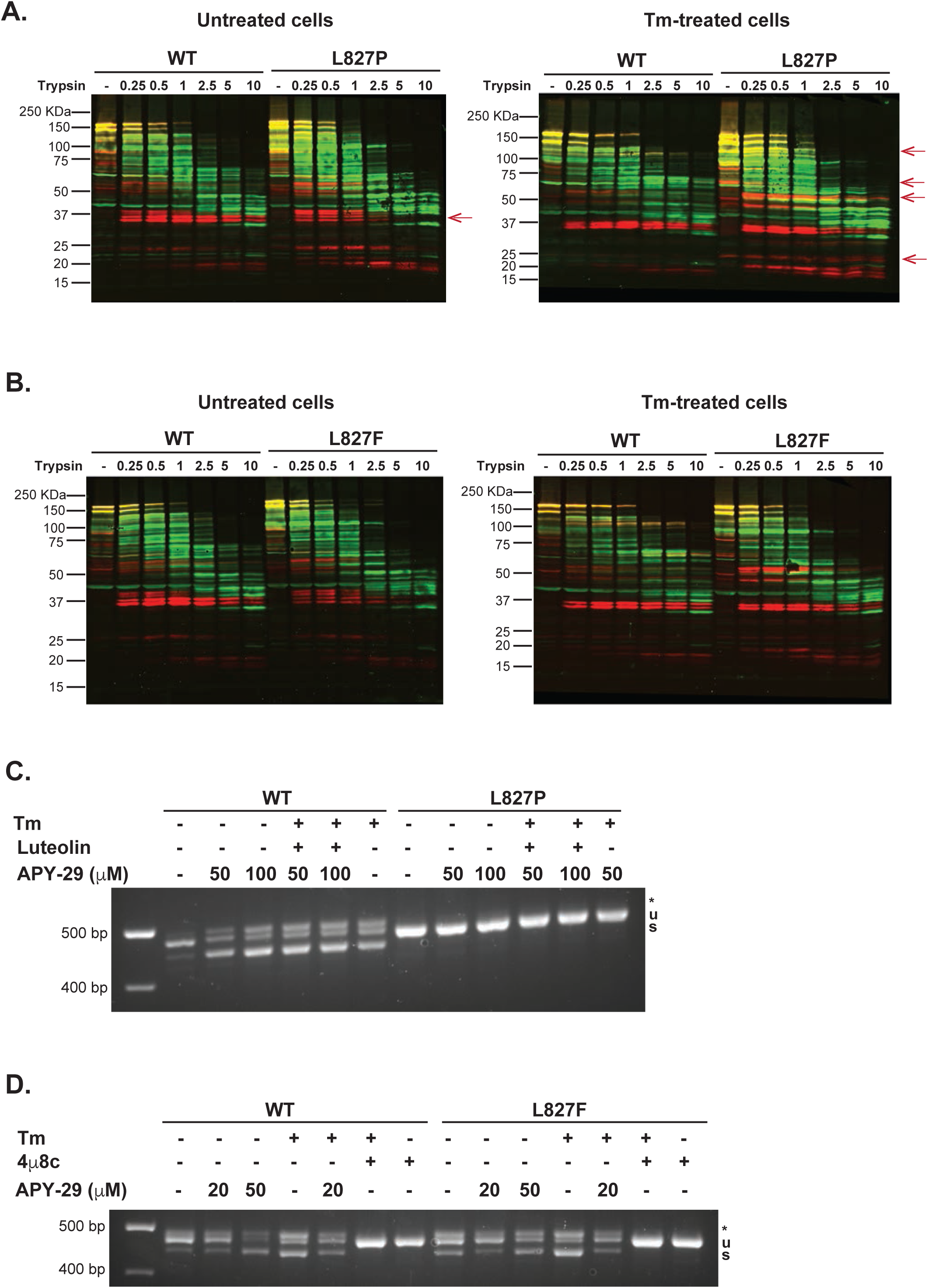
L827P and L827F have conformations distinct from WT IRE1α. **A.** WT and L827P IRE1GFP have different conformations. HAP1KO IRE1GFP WT or L827P cells untreated (left panel) or treated with Tm 4 μg/ml for 4 hours (right panel) were lysed and subjected to the indicated range of trypsin concentrations (μg/ml) for 30 min on ice. Western blot analysis was performed and the membranes probed with anti-GFP (in green) and anti-HA (in red). Yellow bands contain both tags. The GFP domain is 27.5 kDa. The red arrows point to trypsin-induced fragments that differ between WT and L827P. **B.** L827F conformation differs from WT IRE1GFP. The same procedure described in A was performed on either WT or L827F IRE1GFP. **C.** The L827P mutant is not activated by APY-29. HAP1KO expressing IRE1GFP WT or L827P were treated with the indicated concentrations of APY-29, luteolin (50 μM) or Tm (4 μg/ml) for 2 hrs. RT-PCR to detect XBP1 splicing was performed and run on agarose gel. **D.** The L827F mutant is sensitive to APY-29. HAP1KO expressing IRE1GFP WT or L827F were treated with the indicated concentrations of APY-29, Tm (4 μg/ml) or 4μ8c (16 μM) for 2 hrs. XBP1 splicing was detected as above.

A second approach that suggests that mutations in the interdomain helix lead to different conformations utilized the type I allosteric inhibitor of the kinase domain, APY-29 [20, 37]. It provides a bypass activation mode for IRE1α based on allosteric change of conformation of the kinase domain. Paradoxically, APY-29 activates the XBP1 splicing activity in cells at concentrations above 20 μM, even without the stressor Tm (Fig. 7C, D), converting IRE1α to an RNase-active, kinase-inactive conformation [20] with a higher oligomeric state. The L827P mutant is refractive to APY-29 treatment (Fig. 7C), while L827F responds to APY-29 (Fig. 7D). The different behaviors suggest that despite its conformational similarity to L827F (Fig. 7A-B), L827P locks IRE1α in a non-responsive conformation. More importantly, the responsiveness of L827F to APY-29 shows that the nature of residue 827 determines whether the RNase domain is activated with an ATP-competitive inhibitor (see [34]), even though it is distal to both the binding site and the catalytic center and has not been implicated in the pathway of activation.

The trypsin sensitivity assay and the response to allosteric inhibitors suggest that Helix αK can impart distinct conformations on IRE1α, some of which can be manipulated using allosteric kinase inhibitors. The bypass activation by luteolin, to which L827P is also refractive (Fig. 1B, 7C), may operate via the same conformation of the interdomain helix αK that mediates the APY-29 bypass.

### The L827P mutation reduces the ability of HAP1 cells to cope with ER stress

Chronic activation of IRE1α is known to be pro-apoptotic [38], and many enzymatically inactive IRE1α mutants are deficient or even dominant-negative for ER stress-mediated apoptosis [20, 39]. In particular, a P830L mutation near residue L827 causes loss of the enzymatic activities and yet when overexpressed has only subtle inhibitory effect on XBP1 splicing by the WT IRE1α [36] and does not slow cell growth [20]. Therefore, we asked whether the dominant-negative inhibition of endogenous WT IRE1α by L827P has important consequences for ER stress resistance. To avoid the confounding effects of chronic stress, we used a colony formation assay, where cells were exposed to a brief period of ER stress, washed, re-plated in normal growth medium and then evaluated for growth 6 days later. As shown in Fig. 8A, a mild dose of Tm (0.5 μg/ml for 4 hr) was sufficient to inhibit HAP1 colony growth. Expression of the L827P IRE1GFP protein in these cells caused further dramatic decrease in cell survival in response to the Tm stress (Fig. 8B).

We conclude that unlike previously reported inactivating mutations, L827P reduces the cells’ ability to cope with ER stress, likely due to its ability to suppress the cytoprotective WT IRE1α activity *in trans*.

**Figure 8.**
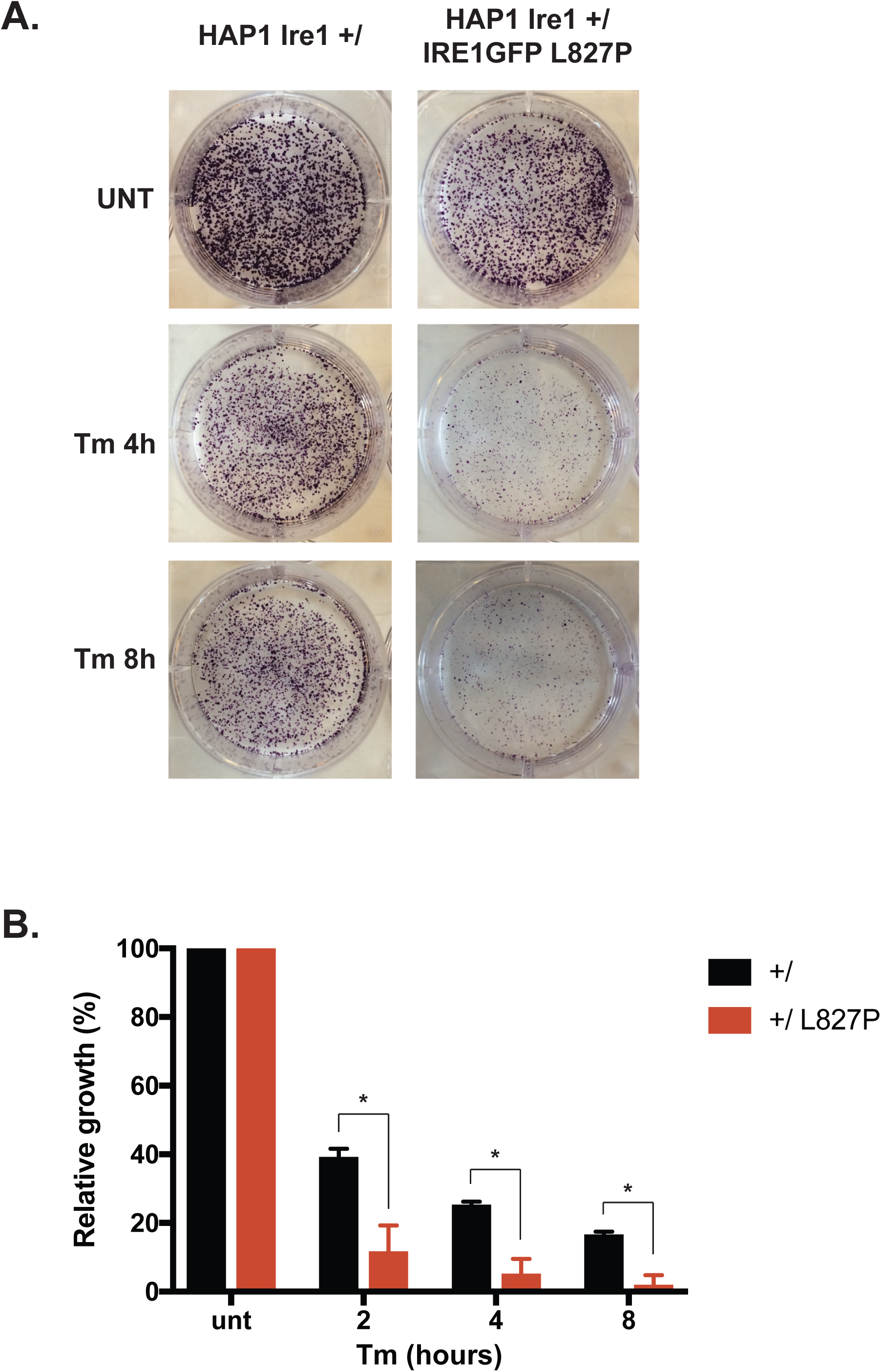
The L827P mutant renders cells expressing endogenous IRE1α more sensitive to ER stress. **A.** The L827P mutant increases the sensitivity of leukemic HAP1 cells to ER stress. Parental HAP1 cells expressing endogenous, WT IRE1α and HAP1 cells expressing both endogenous WT IRE1α and L827P IRE1GFP were induced with dox as above and then treated with 0.5 μg/ml Tm for the indicated time, after which the cells were washed and allowed to grow for 6 days. Colonies that developed by that time were stained with crystal violet. **B.** Quantification of cell growth. The crystals from the plates shown in A were dissolved in 2% SDS and the OD570 of the solutions were quantified. *, p<0.05.

## Discussion

The mutational analysis in this paper defines an intramolecular relay through which the kinase domain of IRE1α communicates to the RNase domain. The trans-auto-phosphorylation that occurs when IRE1α dimerizes is relayed through helix αK residues L827-W833, whose sequence determines the quality of conformational change relayed to the RNase active site (data here and in [36]). Mutations in this helix affect all the measurable activities of IRE1α – XBP1 splicing, RIDD activity, homo- and hetero-oligomerization, and thereby impact the response to ER stress. Even though helix αK is in the cytosolic portion of IRE1α and not in the luminal domain nor in the transmembrane segment, which are each known to sense ER stress, it is necessary for proper activation of IRE1α.

Mutations in helix αK cause IRE1α to assume distinct conformations in response to ER stress, different from WT IRE1α. The most detrimental substitution is L827P, which abolishes the autophosphorylation of the sensor, its ability to splice XBP1, to cleave RIDD transcripts and to cluster in response to ER stress. L827P is also refractive to the bypass activation of IRE1α flavonoids [16] and by allosteric kinase inhibitors [20]. L827 is adjacent to the previously described loss-of-function mutations P830L and W833A [36]. We show that the three amino acids and their interacting residues form an evolutionary conserved, functionally important signaling relay between the kinase and RNase domains. Importantly, even the most severe mutant, L827P, is not a misfolded protein, as it is able to dimerize with wild type IRE1α and co-cluster with it (e.g. Fig. 2 and 3).

Other substitutions of L827 and P830 (L827R, L827F, L827Q, P830A) yield intermediate phenotypes, consistent with the predicted effects of these substitutions on the structure of IRE1α. The less severe substitutions distinguish the inhibition of clustering from the inhibition of RNase activity, and reinforce previous data showing that ER stress-responsive clustering of IRE1α is a distinct manifestation of activation [23].

The severe phenotype of the L827P and the milder phenotype of other substitutions suggest an effect due to altered backbone conformation of this connecting segment. Based on the crystal structures, L827P is predicted to: a) alter the backbone conformation of helix αK between the kinase domain and the RNase domain (Fig. 4); b) create steric clashes with residues A823, H825 and V826 at the C-terminal end of the kinase domain; c) disrupt the proximity of L827 to residues T673 and A677 in helix αE of the kinase domain. It stands to reason that such local effects of the interdomain helix αK mutations would alter the positioning of the RNase domain relative to the kinase domain, perhaps determining the face-to-face or back-to-back orientation of the IRE1α monomers and therefore affecting the activation of the RNase domain [18, 19]. However, an important caveat is that the crystal structures of wild type human IRE1α in the apo form (5HGI [34]), the non-phosphorylated form (3P23 [17]) and phosphorylated pIRE1α (4YCZ and 4YZ9 [40]), all show the helix αK at essentially super-imposable conformations. Thus, the conformational changes that our experimental data imply have not yet been captured in the X ray structures of IRE1α. Ongoing molecular dynamics simulations may provide structural insights that explain the different phenotypes of mutants in the interdomain helix αK.

The transmission of conformational changes along the axis of the IRE1α molecule had long been considered to require auto-phosphorylation at several sites [41], but it was later shown that the phosphoryl transfer *per se* is not essential [18-20]. Instead, the activated kinase initiates a conformational change which is transmitted to the RNase domain. The phosphorylation requirement can be bypassed by mutations like I642G [18] or by using effector small molecules like APY-29 [20], both changing the kinase active site itself. The kinase to RNase conformational change is not well-defined at present and this work suggests that it involves helix αK. None of the known phosphorylation sites [41] are in or near this helix and yet L827P and P830L are not phosphorylated and L827F is phosphorylated in a non-canonical manner. Therefore, it seems that the interdomain helix αK can adopt several intermediate conformations needed to activate the RNase domain, and only some of them are captured in the crystal structures. Since there are conformations that are compatible with enzymatic activity but incompatible with oligomerization, the kinase-RNase domain interface behaves as a tuner that can allow distinct outcomes of activating the IRE1α sensor.

There are subtle but potentially important differences among loss-of-function mutants in helix αK. First, unlike L827P, P830L and W833A were not shown to physically interact with wild type IRE1α; this likely explain why neither of them acted in dominant-negative fashion when IRE1α activation was assayed by phosphorylation or oligomerization [20]. Second, P830L and W833A IRE1α were degraded more rapidly than wild type [36], while the turnover of L827P mutant is unchanged. Our analysis of the predicted free energies of the various substitutions (Table 2) suggests that P830L is a less severe mutation than L827P, so perhaps the phenotypic differences reflect either distinct cell expression environments used in this work and in Xue et al. [36], and/or differential affinity of P830L monomers to WT monomers. A third interesting difference between P830L and L827P is their responsiveness to the kinase inhibitor APY-29: even though both of these mutants are RNase inactive, the phosphorylation activity of P830L is *decreased* by APY-29 and its oligomerization is *increased* [20], whereas L827P is refractive to this drug treatment. This differential responsiveness of mutant IRE1α suggests that their conformations are different.

The dominant-negative nature of L827P is evident not only when over-expressed in fibroblasts, but also in relevant lymphoid cell lines where the mutants are expressed at nearly normal level. This dominant-negative phenotype leads to hyper-sensitivity to chemical ER stress, underscoring the importance of the IRE1α stress sensing pathway for survival of leukemias and lymphomas ([42-45]). So far, no dominant-negative IRE1α has been described in patients or animals, but the mutants described here suggest that there are binding sites outside the kinase or RNase active sites that can be targeted to mimic the pro-apoptotic conformation seen in the L827P mutant. Moreover, the IRE1α kinase active site is structurally similar to other kinases, and therefore targeting a connecting segment whose function depends on kinase activity might be a new selective strategy.

## Acknowledgements

We thank Mr. S. Chomistek and Ms. B. Davies for excellent technical help, Dr. S. Haroon for insightful discussions and Dr. Janis K. Burkhardt for critical reading of this manuscript. We thank Drs. Nathan Roy and Ed Williamson for help with microscopy.

This work was funded by NIH grants AG18001 and GM077480 to Y.A. D. E. was supported by a postdoctoral fellowship from the Italian-American Cancer Foundation. D.R. was supported by NIH training grant 5 T32 HL 7954-18. D.B. was supported by NIH grant R21 AI32828. C.-H.A.T. and C.-C.A.H. were supported by NIH/NCI grant R01CA163910.

## Authors contributions

D.R., S.T., I.L., M.Y., D.E., J.V., S.B. designed and performed experiments.J.P. and A.P. purified the SubAB toxins, C.-H.A.T. and C.-C.A.H provided cells and critical reagents, H.F. and D.B. provided the *in silico* analyses and wrote software, D.R., T.G. and Y.A. wrote the manuscript.

