## Supplementary Figures for "The interdomain helix between the kinase and RNase domains of IRE1α transmits the conformational change that underlies ER stress-induced activation"

**Supplementary Figure Legends**

**Suppl. Figure 1. IRE1α L827P is not phosphorylated in response to ER stress while the L827F mutant is.**

HAP1KO IRE1GFP WT, L827F or L827P were treated with Tm (4 μg/ml) or SubAB (0.3 μg/ml) for 2 hr. The cells were lysed, subjected to phosphatase treatment for 30 minutes at 30°C where indicated and proteins were analyzed by western blot. Arrow: full-length IRE1GFP; §: lower molecular weight species; arrowhead: phosphor-IRE1GFP S729.

**Suppl. Figure 2. Expression levels of exogenous IRE1GFP in multiple myeloma Kms11.**

Expression levels of WT and L827P IRE1GFP proteins in Kms11 cells. Kms11 parental cells or expressing WT or L827P IRE1GFP were induced with dox for 16 hr. Cells were lysed and protein were subjected to western blot analysis. Arrow: full-length IRE1GFP; arrowhead: endogenous IRE1α.

**Suppl. Figure 3. IRE1α L827P does not affect endogenous PERK activity.**

HAP1 cells expressing endogenous IRE1α were complemented with L827P IRE1GFP and exposed to different amounts of dox for 16 hr where indicated. Cells were then treated with Tm (4 μg/ml) for 4 hr, RNA was extracted and relative CHOP mRNA expression was assayed using RT-qPCR quantitation. Data are expressed as the relative abundance of CHOP under each condition relative to the abundance of the unaffected ribosomal gene Rpl19.

**Suppl. Figure 4. L827Q have mild differences with WT IRE1α for ER-stress induced clustering and conformational changes.**

**A.** The ER-stress induced L827Q clusters show some differences with WT ones. HAP1KO IRE1GFP WT or L827Q cells were induces with dox and treated with Tm (4μg/ml) for 4 hr. Images were taken and analyzed using a home-made cluster analyzer for ImageJ. Several parameters were taken into consideration and plotted.

**B.** WT and L827Q IRE1α have similar conformation. HAP1 KO IRE1GFP WT or L827Q cells untreated or treated with Tm 4 μg/ml for 4 hours were lysed and subjected to the indicated range of trypsin concentrations (0-10 μg/ml) for 30 min on ice. Western blot analysis was performed and the membranes probed with anti-GFP (in green) and anti-HA (in red). Yellow bands contain both tags.

**Suppl. Figure 5.** **The stability of mutants L827P, L827F is similar to that of WT IREα.**

Cycloheximide chases (10 μg/ml over 24 hr) of HAP1KO cells expressing WT IRE1GFP, L827P or L827F. Cells were lysed and proteins analyzed by western blot. Arrow indicates full length IRE1GFP; §: lower molecular species which appear to be cycloheximide sensitive. Anti-CD147 antibody was used as a positive control for cycloheximide treatment. Mat.: mature form of CD147; CG: core glycosylated CD147. 14.3.3: housekeeping protein.

**Suppl. Fig. 1**

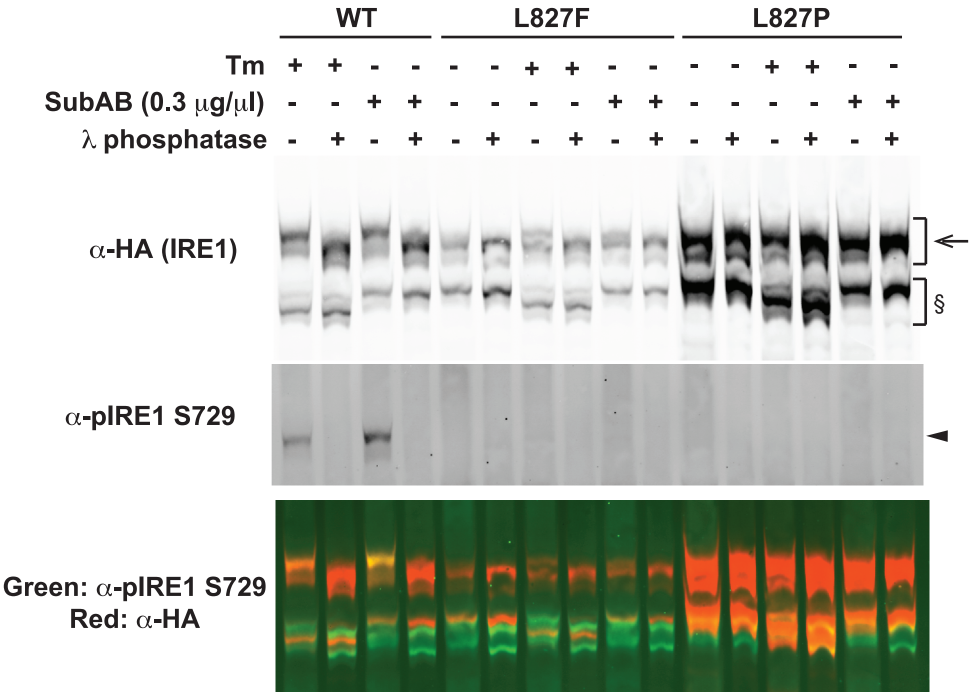
WT
L827F
L827P
Tm
SubAB (0.3 μg/μl)
λ phosphatase
α-HA (IRE1)
§
α-pIRE1 S729
Green: α-pIRE1 S729
Red: α-HA

### Suppl. Fig. 2

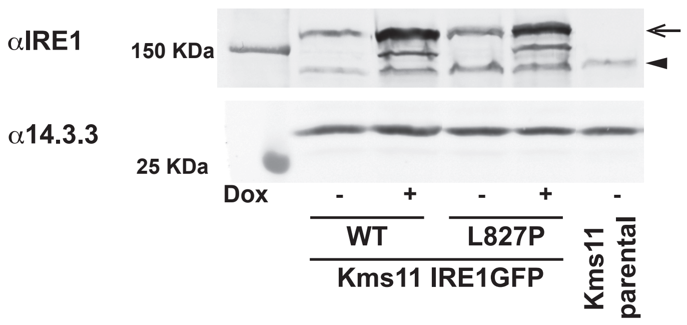
αIRE1
150 KDa
α14.3.3
25 KDa
Dox
-
+
-
+
-
WT
L827P
Kms11 IRE1GFP
Kms11
parental

### Suppl. Fig. 3

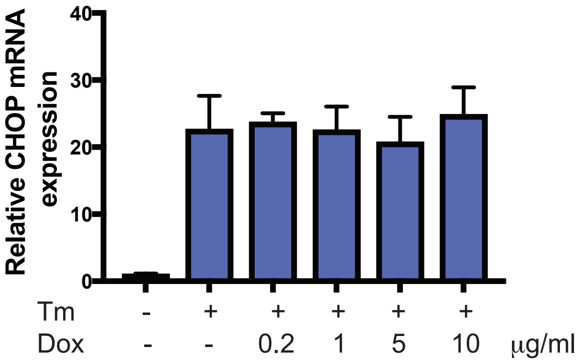

Relative CHOP mRNA expression
40
30
20
10
0
Tm - + + + + +
Dox - - 0.2 1 5 10 μg/ml

Suppl. Fig. 4

A.

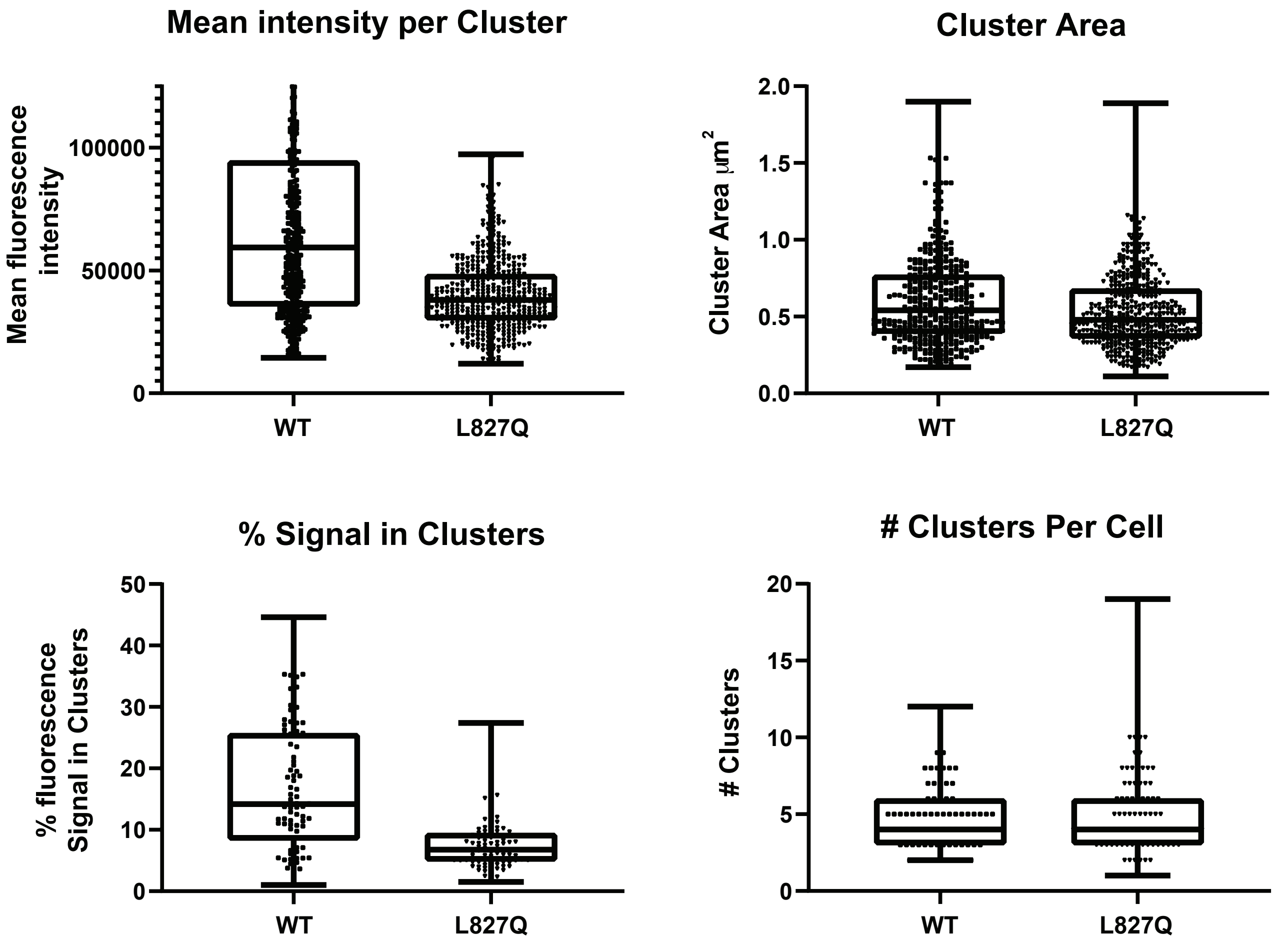


B.

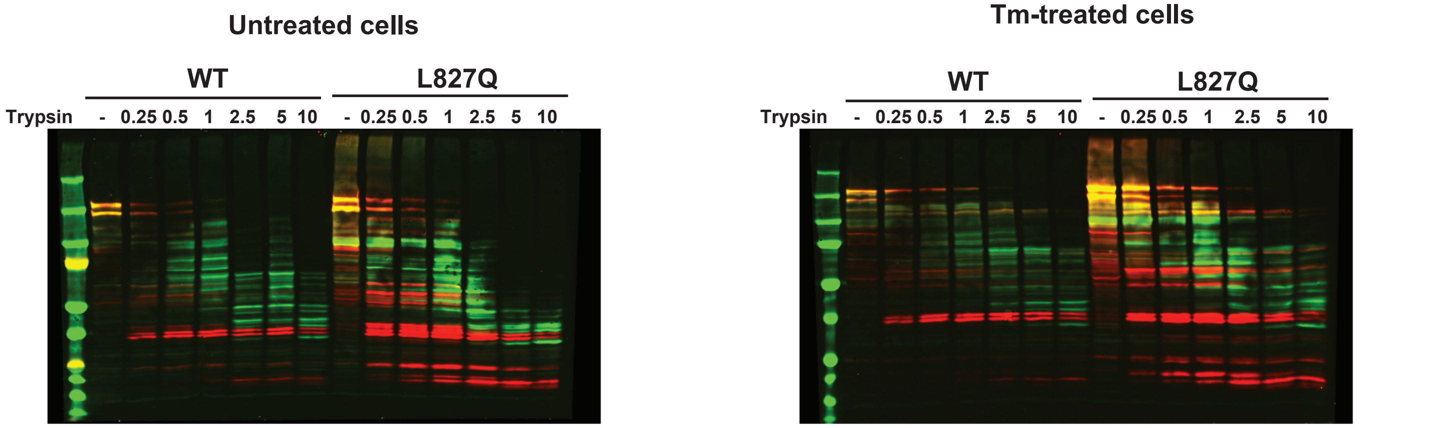

Suppl. Fig. 5

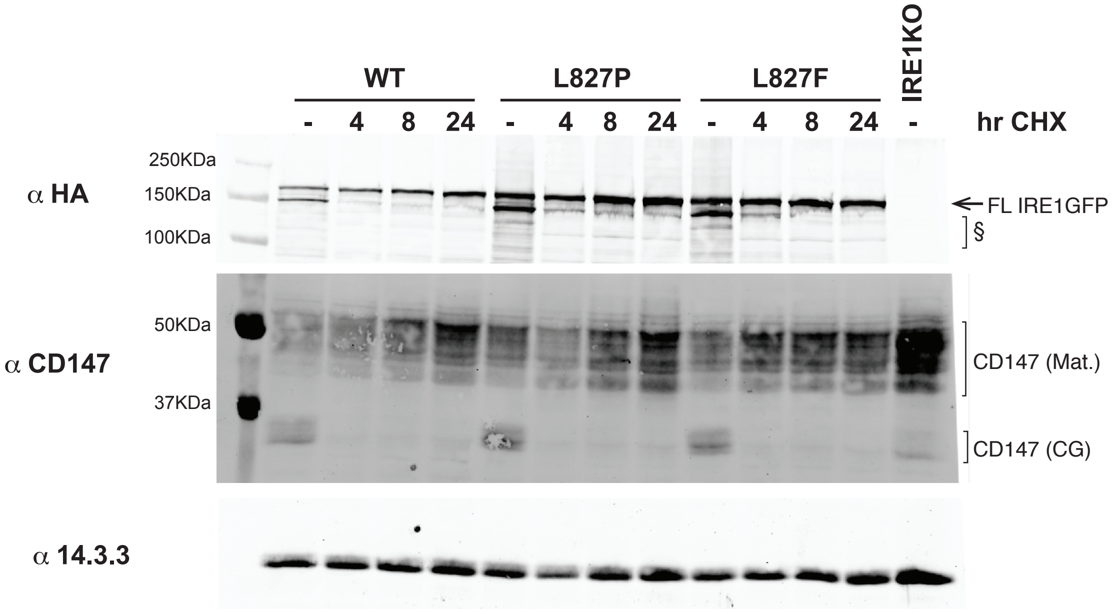
WT
L827P
L827F
IRE1KO
- 4 8 24 - 4 8 24 - 4 8 24 -
hr CHX
α HA
250KDa
150KDa
100KDa
FL IRE1GFP
§
α CD147
50KDa
37KDa
CD147 (Mat.)
CD147 (CG)
α 14.3.3
